# Phenotypic, ecological and genomic divergence support independent evolutionary trajectories in the Meganesian masked owls (*Tyto novaehollandiae*)

**DOI:** 10.64898/2026.11.20.767495

**Authors:** Tristan Cumer, Angélica Pulido, Anne-lyse Ducrest, Céline Simon, Alexandre Roulin, Jérôme Goudet

## Abstract

Understanding how phenotypic, ecological, and genomic divergence accumulates during lineage differentiation remains a central challenge in evolutionary biology. Insular populations provide valuable opportunities to investigate this process because geographic isolation can restrict gene flow and promote divergence. Mainland Australia and Tasmania form an ideal natural system for studying these mechanisms. The Australian masked owl (*Tyto novaehollandiae*) occurs across diverse habitats on mainland Australia and Tasmania. Among its recognized subspecies, the Tasmanian masked owl (*T. n. castanops*) is notable for its larger body size and distinctive plumage compared with Australian *T. n. novaehollandiae*. Here, we combined phenotypic measurements, ecological niche analyses, and whole-genome sequencing to assess divergence between Australian and Tasmanian masked owls.

Tasmanian masked owls were consistently larger and darker reddish in coloration than mainland individuals and occupied a distinct climatic niche. These phenotypic and ecological differences were matched by genomic structure separating individuals by geographic origin. Despite this differentiation, genome-wide genetic divergence remained moderate (F_ST_ = 0.086; D_XY_ = 0.087%), indicating that phenotypic and ecological divergence accumulated more rapidly than genomic differentiation. Genomic analyses also revealed higher inbreeding levels in Tasmanian populations based on runs of homozygosity (F_ROH_). This pattern was driven mainly by short tracts, suggesting long-term isolation rather than recent inbreeding. Genome scans identified several differentiated regions, although no significant enrichment of functional pathways was detected. Overall, our results support the recognition of Australian and Tasmanian masked owls as independently evolving lineages and demonstrate that substantial phenotypic and ecological differentiation can arise before pronounced genome-wide divergence accumulates.

**Significance statement:** Geographically isolated populations often evolve distinct traits, but determining whether they represent independently evolving lineages remains challenging because phenotypic, ecological, and genetic divergence do not always accumulate at the same pace. By combining morphology, climate niche analyses, and whole-genome data, we show that Tasmanian and mainland Australian masked owls differ consistently across all three dimensions despite only moderate genome-wide genetic divergence. These findings demonstrate that substantial evolutionary differentiation can arise rapidly in isolated populations before deep genomic divergence evolves, providing new insights into the early stages of lineage formation.

## Introduction

Understanding how lineages diverge remains a central goal of evolutionary biology. Population divergence is a gradual process during which phenotypic, ecological and genetic differences accumulate through the combined effects of drift, selection and demographic history (Mayr 1963; Futuyma et al. 2023). However, these different components of divergence need not proceed at the same rate. Populations may exhibit substantial morphological differentiation despite limited genomic divergence, whereas strong genetic structure can arise in the absence of obvious phenotypic differences (Nosil 2012; Roux et al. 2016; Zhang and Wiens 2026). Consequently, determining whether geographically isolated populations represent independently evolving lineages or simply differentiated populations remains a recurring challenge. Increasingly, integrative approaches combining phenotypic, ecological and genomic data have been advocated as a robust framework for evaluating lineage divergence and species limits (De Queiroz 2007; Padial et al. 2010; Edwards et al. 2016).

Islands provide particularly valuable opportunities to investigate the processes underlying lineage divergence. Their isolation naturally restricts gene flow and promotes the accumulation of independent evolutionary changes through genetic drift and local adaptation (Macarthur and Wilson 1967; Losos and Ricklefs 2009). Additionally, islands frequently present environmental conditions that differ from those of nearby mainland regions, creating opportunities for ecological divergence (Carson 1974; Warren et al. 2015). The interaction between geographic isolation and ecological differentiation has long been recognised as a major driver of diversification, leading to the evolution of endemic forms across numerous taxa (Grant and Grant 1997; Schluter 2009). Integrating ecological and genomic evidence can therefore provide important insights into whether geographically isolated populations are following independent evolutionary trajectories.

Mainland Australia and Tasmania represent a natural system in which to study these questions. Although currently separated by the Bass Strait, repeated fluctuations in sea level during the Pleistocene periodically connected and isolated Tasmania from the mainland (Lambeck 2004; Byrne et al. 2008). Consequently, Tasmanian vertebrates display heterogeneous phylogeographic patterns, with levels of divergence from mainland populations ranging from minimal differentiation to the persistence of distinct evolutionary lineages (Burridge et al. 2013; Kreger et al. 2020). Determining where lineages lie along this continuum is of particular importance in taxa for which taxonomic assessments have historically relied on morphological evidence alone.

The Australian masked owl (*Tyto novaehollandiae*) is a large nocturnal raptor distributed across a diversity of habitats in mainland Australia and Tasmania. Among its described subspecies, the Tasmanian masked owl, *T. n. castanops*, has attracted attention because of its distinctive morphology, including larger body size and darker plumage traits relative to mainland *T. n. novaehollandiae* (Higgins et al. 2006). These forms also occupy contrasting environments, with Tasmanian populations occurring primarily in cooler and wetter ecosystems than those inhabited by mainland populations (Higgins et al. 2006). While such phenotypic and ecological differences suggest a history of independent evolution, it is yet unclear whether *T. n. castanops* and *T. n. novaehollandiae* represent diverging evolutionary lineages exhibiting concordant genomic differentiation or geneticaly similar populations whose differences are largely restricted to adaptive phenotypic traits. A pattern of divergence at multiple biological dimensions is expected when populations have followed largely independent evolutionary trajectories over evolutionary timescales (De Queiroz 2007; Edwards et al. 2016).

Here, we combine phenotypic measurements, ecological analyses and whole-genome data derived from historical museum specimens to investigate divergence between *T. n. castanops* and *T. n. novaehollandiae*. Specifically, we ask whether these taxa exhibit coherent phenotypic, ecological and genomic differentiation consistent with independent evolutionary trajectories. We quantify phenotypic and ecological differentiation between Tasmanian and mainland Australian populations. We characterise patterns of genomic structure and diversity to assess the extent of evolutionary independence between the two taxa. Finally, we examine patterns of inbreeding and genomic differentiation to evaluate the potential roles of geographic isolation and local adaptation in shaping divergence across environments. By integrating multiple complementary lines of evidence, we evaluate the extent to which these distintive forms of the masked owls are following divergent evolutionary trajectories and provide insight into how phenotypic, ecological and genomic divergence accumulate during the early stages of lineage differentiation.

## Results

Phenotypic measurements from museum specimens revealed marked differentiation between the two taxa. Overall, Tasmanian masked owls (*T. n. castanops*) were larger and darker than mainland Australian individuals (*T. n. novaehollandiae*). Individuals from Tasmania exhibited longer wings (p < 2.2 × 10□¹□), larger bills (p = 1.93 × 10□□), and longer claws (p = 0.010; Fig. 1A to 1C, Supplementary Tables S1–S4). In addition, Tasmanian owls displayed darker plumage coloration on both the breast and the belly (both p < 2.2 × 10□¹□; Fig. 1D and 1E, Supplementary Tables S5and S6). In contrast, the extent of eumelanin spotting (defined as the number of spots) did not differ between taxa, neither on the breast (p = 0.421) nor on the belly (p = 0.060; Supplementary Fig. S1, Supplementary Tables S7 and S8).

**Figure 1.**
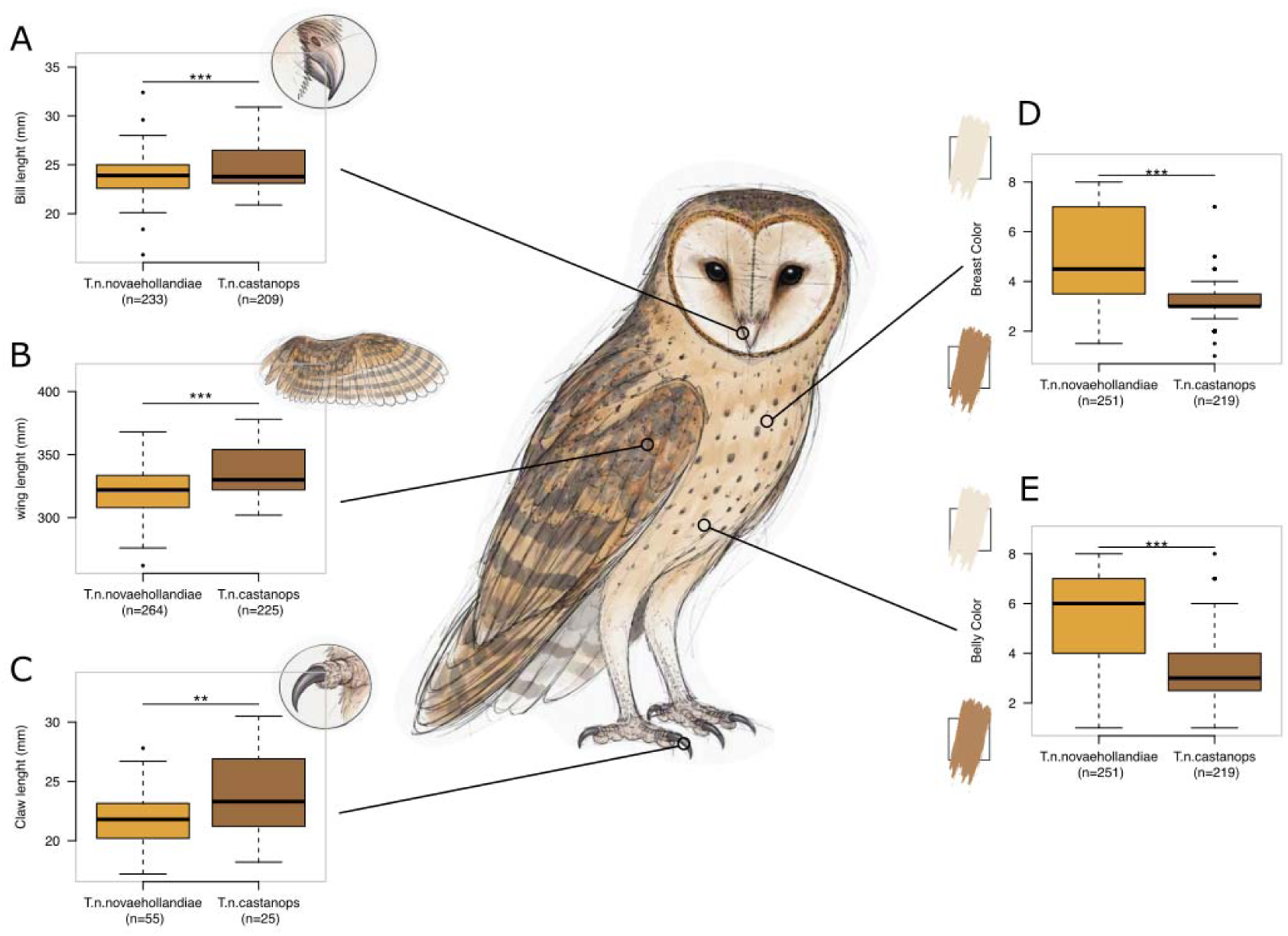
Phenotypic differentiation between the Australian masked owl (*Tyto novaehollandiae novaehollandiae*) and the Tasmanian masked owl (*Tyto novaehollandiae castanops*). Panels A–E respectively show boxplots of bill length, wing length, claw length, breast colour, and belly colour for each population. Colour scores ranged between 1 for darker reddish to 8 for white. Numbers in parentheses below each boxplot indicate sample sizes. Horizontal bars and asterisks indicate the significance of pairwise comparisons between populations (*P* < 0.05, P < 0.01, *P* < 0.001).

We assessed whether Tasmanian birds occupy a distinct ecological niche relative to their mainland counterparts using occurrence records of *Tyto novaehollandiae* across mainland Australia and Tasmania (Fig. 2A). A principal component analysis of climatic variables showed that the first three axes explained 89.3% of the total environmental variation and were therefore retained for subsequent Outlying Mean Index (OMI) analyses (Fig. 2B, Supplementary Fig. S2-S5 and Supplementary Table S9). The first axis was primarily associated with temperature variables (annual mean temperature, maximum temperature of the warmest month and minimum temperature of the coldest month), together with precipitation during the wettest month, and therefore described a broad temperature-moisture gradient. The second axis was mainly driven by isothermality and precipitation during the driest month, reflecting variation in climatic seasonality and aridity. The third axis was dominated by annual temperature range and minimum temperature of the coldest month, corresponding to a gradient of thermal seasonality.

**Figure 2.**
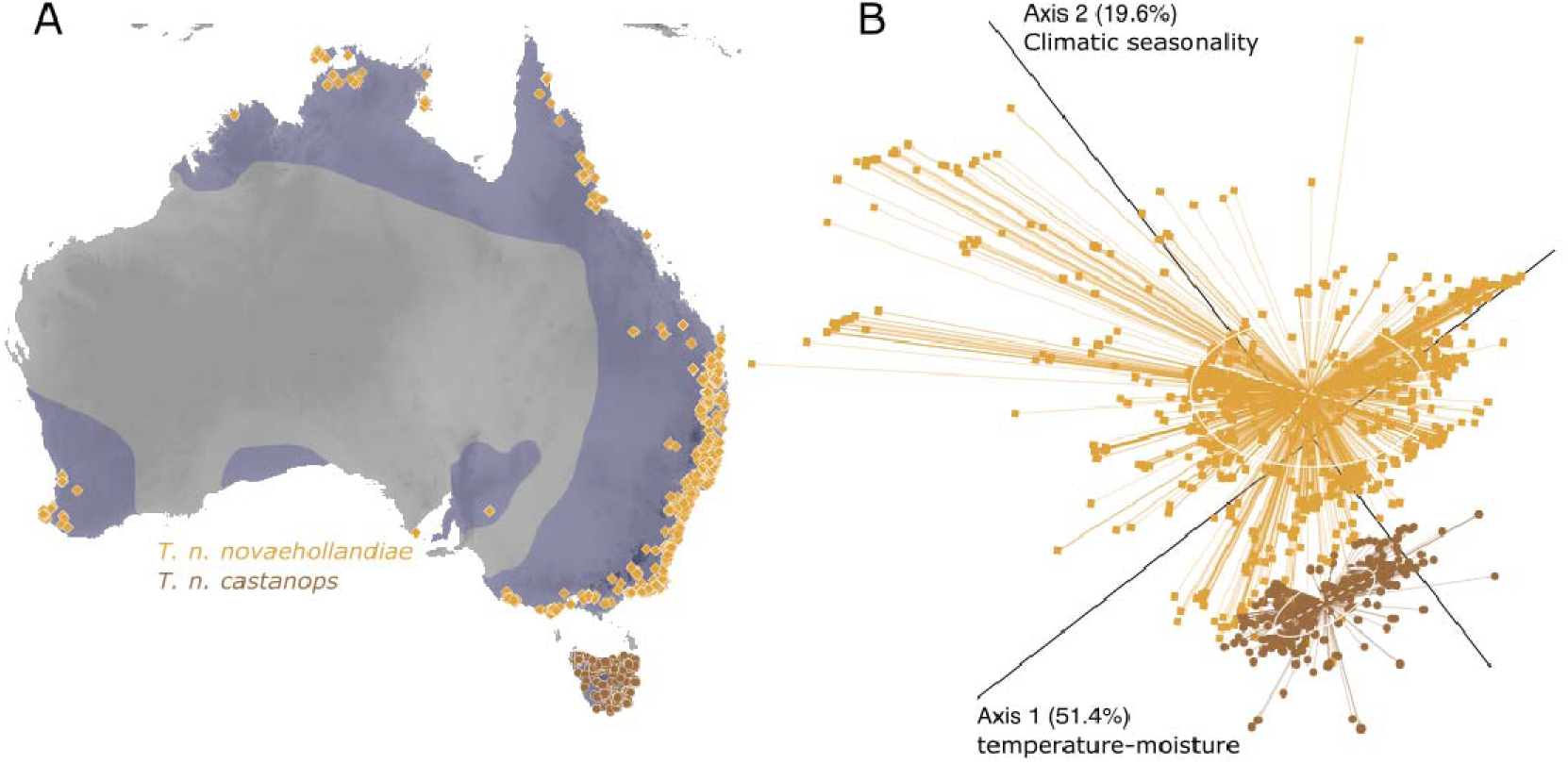
Ecological niches occupied by the Australian masked owl (*Tyto novaehollandiae novaehollandiae*) and the Tasmanian masked owl (*Tyto novaehollandiae castanops*). (A) Geographic distribution of occurrence records used in the analyses (orange: *T. n. novaehollandiae*; brown: *T. n. castanops*). The blue shaded area indicates the species distribution according to the IUCN. (B) Principal component analysis (PCA) of climatic conditions based on 7 bioclimatic variables extracted for all occurrence records shown in panel A. Outlying Mean Index (OMI) analyses revealed significant differentiation in climatic niche occupancy between the two taxa.

The OMI analysis revealed significant ecological differentiation between Tasmanian and mainland Australian populations. Both populations occupied a non-random subset of the climatic conditions available within their respective regions, as indicated by significant marginality values (mainland Australia: OMI = 0.52, p ≤ 0.001; Tasmania: OMI = 7.08, p ≤ 0.001 – Supplementary Table 10). The overall marginality across taxa was also highly significant (OMI.mean = 3.80, p ≤ 0.001), indicating that the climatic niches occupied by Tasmanian and mainland masked owls were more differentiated than expected by chance. Accordingly, *T. n. castanops* occupied a distinct region of climatic space relative to that of *T. n. novaehollandiae*, consistent with the cooler and more climatically stable environments characterising Tasmania relative to most of mainland Australia (Figure 2B).

Given the broad climatic range occupied by *T. n. novaehollandiae* across mainland Australia, we repeated the OMI analysis using only occurrence records from southwestern Australia and Tasmania to assess whether the observed niche differentiation was driven primarily by the extensive environmental heterogeneity of mainland Australia. This restricted analysis yielded results consistent with those obtained across the species’ full range. Ecological differentiation between Tasmanian and mainland populations remained significant (Supplementary Figs. S6–S8; Supplementary Table S11), indicating that the distinct climatic niche occupied by *T. n. castanops* is not driven by the wider environmental breadth encompassed by mainland populations.

We next investigated whether the phenotypic and ecological differences observed between the two taxa were mirrored at the genomic level. Whole-genome sequencing of 21 individuals (9 *T. n. novaehollandiae* and 12 *T. n. castanops -* Supplementary Fig. S9, Supplementary Table S12) yielded 2’476’074 high-quality autosomal SNPs after filtering. Principal component analysis (PCA) revealed a clear separation between the two taxa along the first principal component, which explained 13.5% of the total genetic variance (Fig. 3A, Supplementary Fig. S10). The second and third components explained approximately 5% of the variance each and primarily captured variation among individuals within populations (namely Australia and Tasmania, Supplementary Fig. S10). Consistent with the PCA, ancestry analyses recovered two well-defined genetic clusters corresponding to *T. n. novaehollandiae* and *T. n. castanops* (Fig. 3B). Structure analysis revealed that the best K was K=1, closely followed by K=2, based on the cross-Entropy criterion (Fig. S11). At k=2, Individuals showed high assignment to their population of origin and little evidence of admixture between Tasmania and mainland Australia. Pairwise kinship estimates were consistent with these results (Supplementary Fig. 3C), with individuals being more closely related to members of their own population than to individuals from the other population (mean kinship within *T. n. novaehollandiae* = 0.082; within *T. n. castanops* = 0.019; between populations = −0.035, Fig. S12).

**Figure 3.**
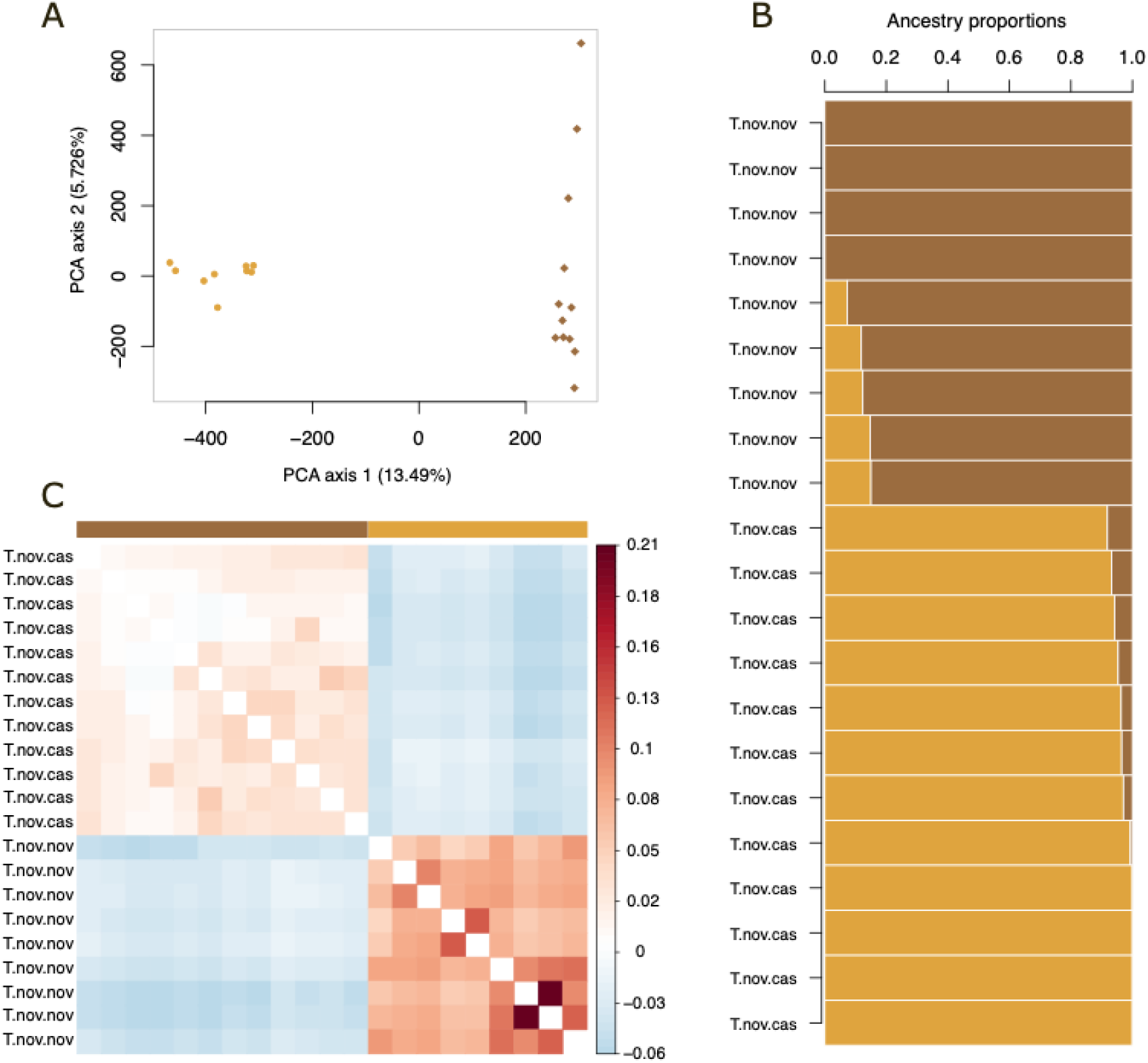
Genetic differentiation between the Australian masked owl (*Tyto novaehollandiae novaehollandiae*) and the Tasmanian masked owl (*Tyto novaehollandiae castanops*). (A) Principal component analysis (PCA) based on genome-wide SNP data. Each point represents an individual (orange: *T. n. novaehollandiae*; brown: *T. n. castanops*). (B) Ancestry analysis assuming *K* = 2 ancestral populations. Each vertical bar represents an individual, and colours indicate the proportional contribution of each inferred ancestral cluster to that individual’s genome. Consistent with the PCA, ancestry analyses revealed clear genetic differentiation between mainland Australian and Tasmanian masked owls.

Given this clear genomic structure, subsequent analyses were conducted by grouping individuals per population. Genome-wide differentiation between populations was small to moderate, with a pairwise F_ST_ of 0.086 (SD = 0.005) and an absolute divergence (D_XY_) of 0.00073 (SD = 2.03e-07). Nucleotide diversity was significantly higher in *T. n. castanops* than in *T. n. novaehollandiae* (π = 0.00083, SD = 1.73e-07 and π = 0.00077, SD = 2.20e-07, p-value < 2.2e-16).

We further characterised genomic inbreeding using a genome wide allele sharing approach (F_S_) combined with a runs-of-homozygosity (ROH) approach (F_ROH_). The two metrics gave contrasting results. The allele sharing approach revealed that the two populations had similar levels of genome wide inbreeding, with Tasmanian *T. n. castanops* displaying higher heterogeneity than mainland *T. n. novaehollandiae* (F_S_= -0.036 (SD = 0.07) and -0.035 (SD = 0.04), respectively, p-value = 0.6; Supplementary Fig. S13). In contrast, Tasmanian *T. n. castanops* exhibited higher levels of genomic inbreeding than mainland *T. n. novaehollandiae* when considering ROH (mean F_ROH_ = 0.13 (var = 0.014) and 0.06 (var = 0.041), respectively; Fig. 4A; Supplementary Fig. S14). This increase in F_ROH_ was primarily driven by an excess of short ROH segments rather than long tracts of homozygosity (Fig. 4B; Supplementary Fig. S15).

**Figure 4.**
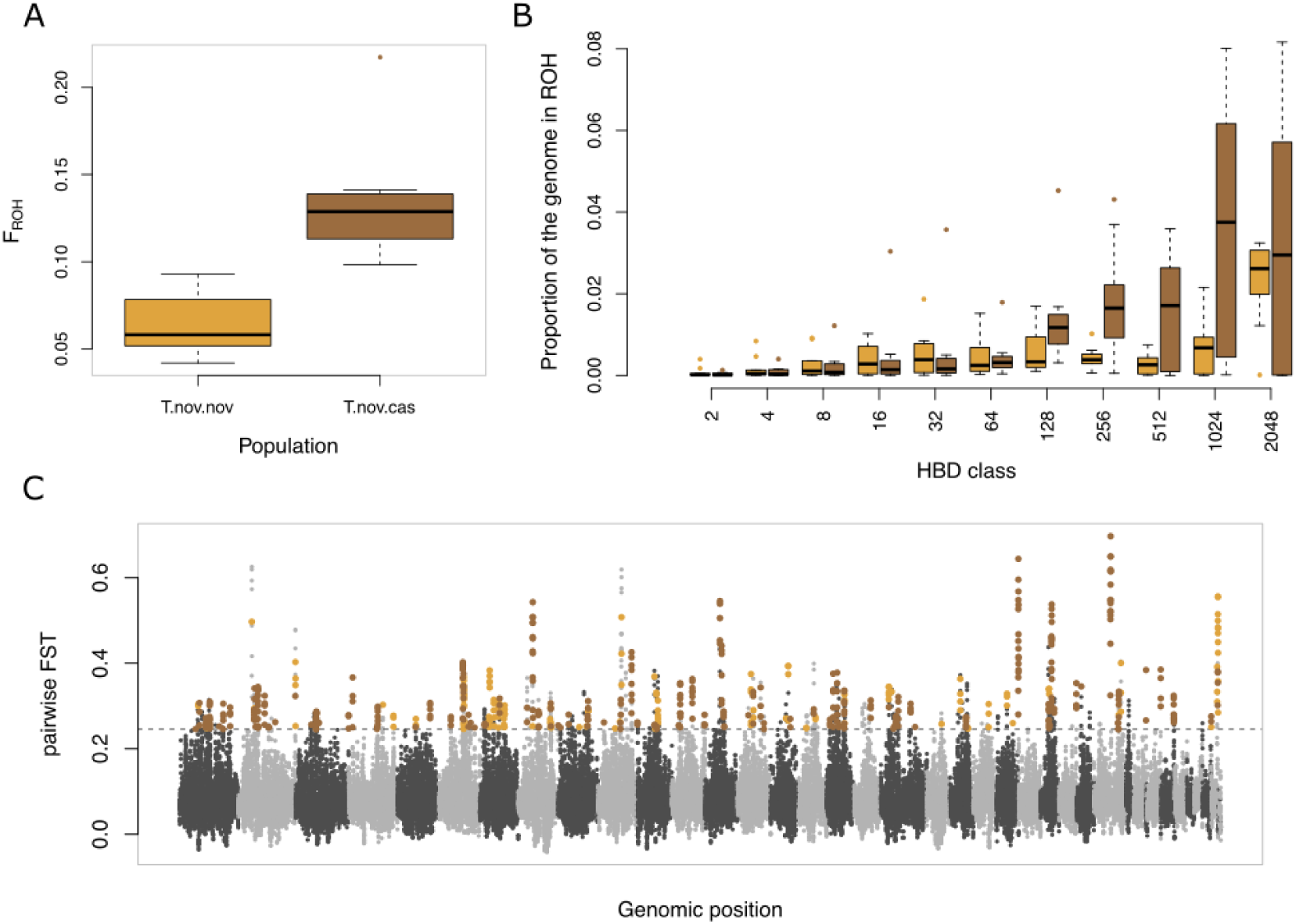
Genome-wide *F*_ST_ comparison between Australian masked owl (*Tyto novaehollandiae novaehollandiae*) and Tasmanian masked owl (*Tyto novaehollandiae castanops*). Each dot represents a 100kbp window. Alternating grey colours denote different chromosomes. Dashed line indicates the 3 SD *F*_ST_ threshold used to identify genomic regions of high differentiation overlaid with *F*_ST_^Tnn^, highlighted in orange and *F*_ST_^Tnc^ in brown.

Considering the moderate genome-wide differentiation between *T. n. novaehollandiae* and *T. n. castanops,* we investigated whether divergence was concentrated in specific genomic regions by combining pairwise F_ST_ and population-specific F_ST_ estimates. This approach identified 13 Mb and 26 Mb of the genome exhibiting both elevated differentiation between taxa and excess relatedness within *T. n. novaehollandiae* and *T. n. castanops*, respectively (Fig. 4C, Supplementary Fig. S16). These regions constitute candidate targets of lineage-specific evolutionary processes. Gene ontology analyses of genes located within these regions (Supplementary Tables S13 and S14) revealed no significant enrichment for biologically relevant pathways. Several candidate (9) loci belonged to the avian β-defensin/gallinacin gene family. Nine candidate loci mapped to the same defensin-related SwissProt orthologue, suggesting local gene duplication or diversification within antimicrobial peptide gene clusters. Together with additional β-defensin candidates (DEFB125, DEFB136), these results point to immunity as a contributor to genomic differentiation.

## Discussion

Divergence is a multifaceted process involving phenotypic, ecological, and genomic changes that do not necessarily accumulate at the same pace. Morphological differentiation may arise through plastic responses to environmental conditions, while genomic divergence can emerge through neutral processes such as geographic isolation and drift. Increasing emphasis has therefore been placed on integrating multiple sources of evidence when evaluating evolutionary divergence and lineage boundaries (De Queiroz 2007; Padial et al. 2010).

By combining phenotypic measurements, ecological niche analyses, and whole-genome data, our study reveals consistent differentiation between *Tyto novaehollandiae novaehollandiae* residing in mainland Australia and *Tyto novaehollandiae castanops* residing in Tasmania. Tasmanian masked owls are larger, darker, and occupy a distinct climatic niche relative to their mainland counterparts. These differences are mirrored by clear genomic structure, with PCA, ancestry analyses, and kinship estimates consistently separating individuals according to their geographic origin. Together, these results indicate that divergence between Australian and Tasmanian masked owls is occurring across multiple biological dimensions.

Despite this concordant differentiation, genome-wide genetic divergence remained moderate (F_ST_ = 0.086). Similar patterns are increasingly recognised in natural populations, where phenotypic and ecological divergence accumulate more rapidly than genome-wide differentiation (Schluter 2009; Nosil 2012; Roux et al. 2016). Moderate genomic differentiation can arise when populations remain relatively recently isolated and continue to retain substantial ancestral variation. Under such circumstances, ecological and phenotypic differentiation may become apparent before extensive divergence accumulates across neutral regions of the genome. This interpretation is consistent with the geological history of the region. Tasmania was repeatedly connected to mainland Australia during Pleistocene glacial periods, with the final inundation of the Bassian land bridge occurring approximately 8,000 years ago following post-glacial sea-level rise (Blom and Alsop 1988). Although sufficient time has elapsed for independent evolutionary trajectories to emerge, this period remains relatively short compared with the timescales often associated with deep genome-wide divergence (Roux et al. 2016). Consequently, the observed pattern of strong phenotypic, ecological, and genomic structuring coupled with moderate F_ST_ is consistent with two lineages undergoing substantial divergence despite their relatively recent geographic isolation.

Island populations are frequently expected to exhibit reduced genetic diversity, elevated inbreeding, and the accumulation of long runs of homozygosity as a consequence of small effective population sizes and prolonged isolation (Frankham 1997; Keller and Waller 2002). If the divergence of *T. n. castanops* had been driven primarily by strong genetic drift following isolation, pronounced genomic erosion and clear signatures of recent inbreeding would therefore be expected.

Our results provide only partial support for elevated inbreeding in the insular population. Although *T. n. castanops* exhibited higher genomic inbreeding than mainland owls when estimated from runs of homozygosity (F_ROH_), genome-wide allele-sharing estimates (F_S_) revealed similar levels of inbreeding in the two populations. This apparent discrepancy is not unexpected, as the two metrics capture different aspects of genomic variation. Allele-sharing approaches (F_S_) primarily reflect genome-wide heterozygosity, whereas F_ROH_ estimates the fraction of the genome contained within homozygous-by-descent segments and therefore more directly reflects realized autozygosity (Kardos et al. 2015; Ceballos et al. 2018). Consequently, populations may display similar levels of overall heterozygosity while differing substantially in the extent and genomic distribution of homozygous-by-descent tracts. Such a pattern is expected when historical isolation and genetic drift increase autozygosity without causing a marked reduction in contemporary genome-wide heterozygosity (Ceballos et al. 2018).

In *T. n. castanops*, the higher F_ROH_ was driven primarily by an excess of short homozygous tracts rather than long ROH. Long ROH are generally associated with recent matings among close relatives, whereas short ROH reflect more ancient demographic processes such as bottlenecks, founder events, and long-term reductions in effective population size (Ceballos et al. 2018; Kardos et al. 2021). The predominance of short ROH therefore suggests that genomic signatures in *T. n. castanops* are primarily the result of long-term isolation and historic demographic events (e.g bootleneck suffered by the founder population) allowing historical autozygosity to accumulate, rather than demographic collapse due to recent consanguinity.

Unexpectedly, nucleotide diversity was significantly lower in mainland Australian owls than in Tasmanian individuals. This pattern contrasts with the classical expectation for island populations, where prolonged isolation is commonly associated with reduced genetic variation (Frankham 1997). First, the mainland sample analysed in this study comprises only nine individuals distributed across a limited portion of the range of *T. n. novaehollandiae* (Table S12), representing only a small portion of the overall distribution of *T. n. novaehollandiae* (Fig. 2). Consequently, our estimates may reflect only a subset of the genetic diversity present across mainland Australia. This interpretation is further supported by the higher pairwise relatedness observed among mainland individuals (Fig. 3C), suggesting increased background genetic similarity within the sampled population. However, this increased relatedness does not appear to reflect elevated levels of inbreeding, as genome-wide inbreeding coefficients (F_S_) were similar between Australian and Tasmanian populations. Together, these results suggest that the lower nucleotide diversity observed in the mainland sample may primarily reflect reduced local genetic diversity and increased background relatedness rather than increased autozygosity.

Second, large island populations can maintain substantial genetic diversity despite long-term geographic isolation (Zhang and Wiens 2026). Such a pattern has recently been documented in European barn owls, where insular populations retained considerable standing genetic variation despite extended demographic separation (Machado et al. 2022; Lavanchy et al. 2024).

While the diversity and ROH patterns are consistent with a history of demographic isolation in Tasmania, the present dataset is not ideally suited for formal demographic reconstruction. Estimating parameters such as historical effective population size (Ne), divergence time, or changes in gene flow typically relies on site-frequency spectrum-(Liu and Fu 2020), haplotype-(Waples 2006), or coalescent-based approaches (Li and Durbin 2011a), which perform best with high-coverage genomic data, accurate genotype calls, phased haplotypes, and reliable ancestral-state polarisation (Li and Durbin 2011b). In contrast, our analyses were based primarily on genotype likelihoods derived from low-to-moderate coverage genomes, an approach well suited for inference of population structure, diversity, and inbreeding but less optimal for detailed demographic modelling. Future studies incorporating more samples, higher-coverage genomes, broader geographic sampling, and explicit demographic analyses could provide quantitative estimates of divergence times and historical changes in Ne, allowing direct tests of the demographic scenarios suggested by our results.

Taken together, these findings indicate that both demographic and geographic isolation has contributed to divergence between Australian and Tasmanian masked owls, but that drift alone is unlikely to fully explain the observed patterns. Historical isolation is evident in the elevated proportion of short ROH and the clear genomic structure separating the two populations, yet the severe loss of genetic diversity expected under a founder bottleneck and subsequent strong and persistent drift is not observed. Thus, while demographic processes have undoubtedly shaped divergence, additional evolutionary mechanisms are likely to have contributed to the marked phenotypic and ecological differentiation observed between the two lineages.

Collectively, our results suggest that Bass Strait has provided sufficient isolation to promote independent evolutionary trajectories while avoiding the severe genomic erosion frequently associated with small island populations. This interpretation is further supported by the moderate genome-wide differentiation and limited evidence of admixture between mainland and Tasmanian owls. Large water bodies are known to represent substantial barriers to dispersal and gene flow in owls, even when geographic distances are relatively modest (Cumer et al. 2022). Bass Strait therefore appears to have acted as an effective barrier to gene flow following its inundation, facilitating genetic and phenotypic divergence between populations on either side of the strait.

At the same time, Tasmania is sufficiently large to support substantial population sizes and maintain considerable standing genetic variation, facilitating adaptive divergence to contrasting environments through natural selection. This combination of prolonged isolation and retained diversity provides conditions under which both neutral and adaptive evolutionary processes can contribute to divergence.

The phenotypic and ecological differences observed between *T. n. novaehollandiae* and *T. n. castanops* may also be associated with lineage-specific responses to contrasting environments. Ecological niche analyses revealed that Tasmanian and Australian populations occupy distinct climatic conditions, suggesting that they have experienced different selective pressures over their period of isolation. Although the specific environmental drivers remain to be identified, the concordance among ecological, phenotypic, and genomic differentiation is consistent with expectations under environmentally mediated divergence (Nosil 2012).

Supporting this interpretation, genome scans identified several genomic regions exhibiting elevated differentiation between taxa together with increased within-population similarity. While these regions may represent targets of lineage-specific selection, alternative explanations involving demographic history, background selection, and linked selection cannot be excluded. Consequently, these results should be viewed as consistent with, rather than definitive evidence for, local adaptation.

Notably, signals of differentiation were distributed across multiple genomic regions rather than concentrated within a small number of loci. Such a pattern is more consistent with polygenic adaptation than with selection acting on one or a few loci of major effect. Polygenic divergence can arise through the differential sorting of standing genetic variation across heterogeneous environments, allowing populations to respond to environmental change without requiring the emergence of new mutations: recent work in barn owls suggests that standing genetic diversity can act as a reservoir of adaptive variation, facilitating evolutionary responses across both spatially and temporally variable environments (Corval, Cumer, et al. 2026). The divergence observed between *T. n. novaehollandiae* and *T. n. castanops* may therefore reflect the gradual sorting of pre-existing genetic variation following exposure to contrasting climatic regimes in mainland Australia and Tasmania. However, the present dataset does not allow us to determine whether the observed phenotypic differentiation exceeds expectations under neutral drift alone. Future work integrating genotype-environment associations, functional annotation of differentiated genomic regions, and phenotypic data will be essential to determine the extent to which neutral and adaptive processes have contributed to divergence between these lineages. Such analyses would provide valuable insight into the mechanisms linking environmental variation, phenotypic differentiation, and genomic change.

Taken together, the phenotypic, ecological, and genomic evidence support the view that Australian and Tasmanian masked owls represent independently evolving lineages. Although genome-wide divergence remains moderate, consistent differentiation across multiple biological dimensions suggests that evolutionary divergence is well underway. The combination of distinct morphology, climatic niche occupancy, genomic structure, and limited evidence of contemporary gene flow indicates that *T. n. castanops* represents a unique component of masked owl evolutionary diversity.

From a conservation perspective, preserving this diversity may be particularly important given the apparent evolutionary distinctiveness of the Tasmanian lineage and the substantial standing genetic variation retained within this population. Maintaining this diversity could enhance the capacity of populations to respond to future environmental change and help preserve adaptive potential. More broadly, our results highlight the value of integrating phenotypic, ecological, and genomic data when evaluating evolutionary divergence and conservation units, particularly in systems where substantial ecological and phenotypic differentiation has evolved despite only moderate genome-wide genetic divergence.

## Conclusion

Our study demonstrates that *T. n. castanops* and *T. n. novaehollandiae* differ consistently in phenotype, ecology, and genomic composition. Although genome-wide differentiation remains moderate, the concordance among these independent lines of evidence suggests that these lineages are in the early stages of evolutionary divergence and have followed largely independent evolutionary trajectories since the isolation of Tasmania from mainland Australia. The genomic patterns observed indicate that historical geographic isolation has played an important role in shaping this divergence, yet the maintenance of substantial genetic variation suggests that drift alone is unlikely to fully explain the phenotypic and ecological differences between lineages.

More broadly, our findings illustrate how substantial phenotypic, ecological, and genomic divergence can accumulate in geographically isolated populations before the emergence of deep genome-wide differentiation. Bass Strait appears to have restricted gene flow sufficiently to promote divergence while the large size of landmasses it separates allowed populations to preserve much of the standing genetic variation within each lineage. This retained diversity may, in turn, have facilitated adaptive responses to the contrasting environmental conditions of Tasmania and mainland Australia. Together, these results support the view that *T. n. castanops* and *T. n. novaehollandiae* represent independently evolving lineages and highlight the importance of integrating phenotypic, ecological, and genomic evidence when evaluating evolutionary divergence in recently isolated populations.

## Material and Method

### Phenotypic data collection

Phenotypic data were collected from 471 masked owl skins (264 *Tyto novaehollandiae novaehollandiae* - Tnn - and 225 *Tyto novaehollandiae castanops* - Tnc) from 41 different museums (see Supplementary Table S1). Sex was not included in the analyses as it was unavailable for most specimens. Reliable sex determination from phenotypes is generally not possible from museum skins in this species unless a brood patch is present, a condition observed only in a few female specimens. Because sex information was unavailable for most individuals, it was not incorporated into subsequent analyses. Phenotypic measurements including, but were not limited to, bill length (mm), claw length (mm), wing length (mm), the degree of reddish colouration of the breast and belly, and the number of spots on the breast and belly, were recorded. The degree of reddish colouration of the breast was categorised on an eight-level scale ranging from 1 (dark red) to 8 (white). This method has been found highly reliable for describing variation in background feather colour in this species given its strong correlation with reflectance in the visible spectrum, as measured by spectrometric procedures (Dreiss and Roulin 2010). Spot number was estimated by counting the number of eumelanic black spots within a 60 × 40 mm (2400 mm²) frame.

### Climatic niche analysis

To assess whether the masked owl populations from mainland Australia and Tasmania occupy different climatic niches, we used the Outlying Mean Index (OMI) approach, a multivariate method developed to investigate niche separation and niche breadth (Dolédec et al. 2000) as implemented in the R package ade4 v.1.7-24. This method measures the distance between the mean habitat conditions used by a species (species centroid), and the mean habitat conditions of the sampling area (origin of the niche hyperspace). Thus, the position of the species will depend on their niche deviation from the reference, a theoretical ubiquitous species that tolerates the most general habitat conditions (i.e., a hypothetical species uniformly distributed among habitat conditions). Observation points for masked owls were compiled from the Global Biodiversity Information Facility (GBIF.org 2025). We kept records only from the landmasses sampled in this study, namely Australia (*T. n. novahollandiae*; n= 2087) and Tasmania (*T. n. castanops;* n= 558).

Climatic variables for the region were extracted from the WorldClim database (Hijmans et al. 2005) at 30 sec resolution (approximately 1 km^2^) using the R package rbioclim (Exposito-Alonso 2017). Redundant variables (correlation > 0.8) were filtered out (Figure S2 and S3). The final analysis was run with the following variables: Annual Mean Temperature (BIO1), Isothermality (BIO3), Max Temperature of Warmest Month (BIO5), Min Temperature of Coldest Month (BIO6), Temperature Annual Range (BIO7), Precipitation of Wettest Month (BIO13) and Precipitation of Driest Month (BIO14).

To avoid inflating the signal with geographically (and potentially ecologically) distant occurrences, we reran the analyses focussing on a subset of observations restricted to Tasmania and the southeastern part of Australia (*T. n. novahollandiae*; n= 2087, *T. n. castanops;* n= 558).

### Whole-genome sequencing, read mapping and SNP calling

30 specimens of masked owls were sampled from museum skins with individuals representative of the two populations (Supplementary Table S12, Figure S9): Mainland Australia (*T. n. novaehollandiae* - Tnn, n = 12) and Tasmania (*T. n. castanops* - Tnc, n = 18). Genomic DNA was extracted using the DNeasy Blood & Tissue kit (Qiagen, Hilden, Germany). Library preparation was performed by Rapid Genomics for Illumina sequencing utilizing their high-throughput workflow with proprietary chemistry. Briefly, DNA is sheared to target a mean fragment length of 300bp, fragments are end-repaired and A tailed, followed by incorporation of unique dual-indexed Illumina adaptors and PCR enrichment. Samples are pooled equimolar and sequenced using 2×150bp on an Illumina Novaseq 6000.

The bioinformatics pipeline used to obtain analysis-ready SNPs from the 30 individuals follows standard procedures. Considering the potential degradation and short length of the DNA fragments retrieved from museum samples, forward and reverse reads were merged when needed using Pear v.0.9.11 (Zhang et al. 2014). Paired-end and merged reads were trimmed independently with Trimommatic v.0.36 (Bolger et al. 2014) and filtering out reads shorter than 70 bp after trimming (parameter MINLEN:70). This procedure yielded to an average of 74,991,190.4 of merged reads per samples (min 68,58,802; max 184,566,651 - Supplementary Table S12). Since no reference genome is available for the focal species (*Tyto novaehollandiae*), all trimmed reads (paired, merged and unpaired) were aligned against the chromosome level reference genome of *Tyto alba* (Bioproject PRJNA1440222) (Corval, Ducrest, et al. 2026). Mapping coverage ranged between 0.68 and 20.87, with a mean of 7.91 (per sample mapping statics are reported in Supplementary Table S12). For downstream analyses we focussed only on individuals with a mean coverage higher than 3X, resulting in a total of 25 samples (Tnn = 9 and Tnc = 16).

Considering the low to medium depth of coverage, we called variants using the likelihood-based approach implemented in the ANGSD framework (Korneliussen et al. 2014). To ensure high-quality variant identification, only uniquely mapped reads were retained, and minimum thresholds of mapping quality (MAPQ ≥ 30) and base quality (Phred score ≥ 20) were applied. Adjustment for excessive mismatches around potential indels was performed using a coefficient of 50 (-C 50), and base alignment quality (BAQ) recalibration was enabled (-baq 2). Tri-allelic sites were excluded from the analysis. Candidate bi-allelic SNPs were identified using a significance threshold of p< 10^-6^ (-snp_pval 1e-6). Sites with a minor allele frequency (MAF) below 1% were discarded (-minMaf 0.01). To limit the influence of missing data, variants were required to be represented in at least 17 individuals (-minInd 17), while a minimum sequencing depth of 3 reads per individual was required for genotype likelihood estimation (-setMinDepthInd 2). Genotype likelihoods were calculated using the SAMtools model (-GL 1). Major and minor alleles were inferred based on predefined allele frequencies (-doMajorMinor 4), allele frequency estimates were obtained using posterior probabilities (-doMaf 3), and posterior genotype probabilities were computed (-doPost 1). Finally, only SNPs located in the regions of the genome where variants can be reliably called were considered (see (Corval, Ducrest, et al. 2026) for details).

To account for uncertainty associated with low-coverage sequencing in downstream analyses, genotype likelihoods were converted into posterior genotype probabilities and genotype dosages (expected allele counts). Both representations retain information on genotype uncertainty and avoid the loss of information associated with hard genotype calling in low-depth sequencing data. This approach follows the general framework advocated for low-coverage sequencing studies, in which population genetic inference is performed using genotype uncertainty rather than discrete genotype calls (Meisner and Albrechtsen 2018). This final dataset was filtered one last time to discard individuals with more than 10% missing data (4 individuals discarded) and SNPs with more than 5% data and a minor allele count lower than 5. The output consisted of filtered set of 2,476,074 filtered SNPs in a total of 21 samples (Tnn=9 and Tnc=12).

### Population structure and genetic diversity

To investigate population structure among our samples, sNMF v.1.2 (Frichot et al. 2014) was run for K = 2–4 in 25 replicates to infer individual clustering and ancestry coefficients. For these analyses, only SNPs genotyped in all individuals were retained (11’016 SNPs). To minimize the influence of physical linkage among markers, as recommended for ancestry analyses, we generated a dataset of approximately independent SNPs by randomly selecting a single SNP from non-overlapping 100-kb windows across the genome, yielding a set of 11,017 SNPs. This dataset was subsequently used for both sNMF and Principal Component Analysis (PCA), implemented in the R package LEA (Frichot and François 2015).

Genome-wide population differentiation (F_ST_), nucleotide diversity (π) and absolute divergence (D_XY_) were estimated using the complete dataset of 2,476,074 SNPs. Pairwise and population-specific F_ST_ values were calculated using the estimators implemented in the R package hierfstat v.0.5.11 (Goudet 2005; Weir and Goudet 2017). D_XY_ was calculated according to (Nei 1987) using a custom script. Confidence intervals for F_ST_, π and D_XY_ were estimated by randomly bootstrapping SNPs 50 times.

### Genetic Relatedness and inbreeding

To characterise patterns of relatedness within and between populations, we estimated pairwise kinship coefficients using the complete dataset of 2,476,074 SNPs. Kinship coefficients were calculated using the allele-sharing estimator implemented in the R package hierfstat v0.5.11 (Goudet 2005; Weir and Goudet 2017), for which relatedness is expressed relative to the mean kinship among all individuals included in the analysis. Pairwise kinship was estimated both across the entire dataset and within each taxon separately to assess patterns of relatedness among and within populations.

Individual inbreeding was estimated using the allele sharing estimate (Goudet et al. 2018b) implemented in the R package hierfstat v0.5.11 (Goudet 2005; Weir and Goudet 2017). We also used the run of homozygosity (ROH) approach implemented in ZooRoH v0.4.1 (Druet and Gautier 2017). ZooRoH uses a hidden Markov model to identify homozygous-by-descent (HBD) genomic segments and infer the proportion of the genome inherited from common ancestors. For each individual, we calculated F_ROH_, defined as the proportion of the genome contained within ROH segments. Because ROH arise when homologous chromosomal segments are inherited from a common ancestor, both their abundance and length provide information about the demographic processes shaping genomic variation. Long ROH typically result from recent mating between related individuals, whereas shorter ROH reflect older shared ancestry associated with historical bottlenecks, founder events, or long-term small effective population sizes. In addition to estimating overall genomic inbreeding, we examined the distribution of ROH lengths among individuals to distinguish between recent and historical sources of inbreeding and to compare demographic patterns between *T. n. novaehollandiae* and *T. n. castanops*.

### Genome scan for excessive differentiation between lineages

To investigate potential genomic signals of differentiation, putatively linked with ecological adaptation to their distinct niches, we contrasted the two lineages (*T. n. novaehollandiae* – Tnn against *T. n. castanops* - Tnc). To do so, we used the pairwise *F*_ST_ approach combined with the population-specific *F*_ST_ (Weir and Goudet 2017; Goudet et al. 2018b). This combination of approaches allowed us to identify genomic regions of high differentiation between populations (based on pairwise *F*_ST_) and classify these regions as putatively under selection in one or the other population (based on the population specific *F*_ST_). To do so, we first calculated the pairwise *F*_ST_ from allele sharing matrices (Weir and Goudet 2017; Goudet et al. 2018a) in sliding windows of 100kbp with 20kbp steps, using hierfstat v0.5.11. We then computed the population specific *F*_ST_ for each population (respectively *F*_ST_^Tnn^ and *F*_ST_^Tnc^) and for each matrix, by calculating (i) the mean allele sharing for pairs of individuals within populations and (ii) the mean allele sharing for pairs of individuals between populations, from which we obtain the population specific *F*_ST_.

For each population, we consequently identified genomic highly differentiated between islands (regions with windows of pairwise *F*_ST_ higher than 3 SD from the mean) and above 1 SD from the mean difference each population’s *F*. Genes in these regions were extracted from the annotation of the *Tyto alba* reference genome. Candidate genes were annotated through homology to SwissProt proteins (Waterhouse et al. 2018). Gene Ontology (GO) enrichment analyses were conducted using the R package clusterProfiler v3.23 (Yu et al. 2012), with all annotated barn owl genes as the background universe. GO annotations were retrieved from Ensembl BioMart based on HGNC gene symbols, and enrichment significance was evaluated using Benjamini-Hochberg correction for multiple testing.

## Supporting information

Spplementary data

## Data Availability Statement

All phenotypic data supporting this study are provided in Table S1. Species occurrence records used for ecological niche analyses were downloaded from the Global Biodiversity Information Facility (GBIF) and are publicly available through the associated GBIF download (https://doi.org/10.15468/dl.gwpumg) and provided in table S9. Raw genomic sequence data will be deposited in the National Center for Biotechnology Information (NCBI) Sequence Read Archive (SRA) prior to publication and will be released upon publication.

## Acknowledgements

We are deeply grateful to the museum curators who generously provided access to their extensive specimen collections. Without their collaboration, the phenotypic and genomic data presented in this study could not have been collected. A complete list of participating institutions is provided in Table S1.

AI Disclosure: The owl illustration included in Figure 1 was generated using Microsoft Copilot (Microsoft Corporation) and was used solely for visual illustration of the study species.

