## Supplementary material for "Phenotypic, ecological and genomic divergence support independent evolutionary trajectories in the Meganesian masked owls (*Tyto novaehollandiae*)"

### Phenotypic differentiation

Table S1 – Phenotypic records from 489 masked owl skins (264 *Tyto novaehollandiae novaehollandiae* and 225 *Tyto novaehollandiae canstenops*) from 41 museums.

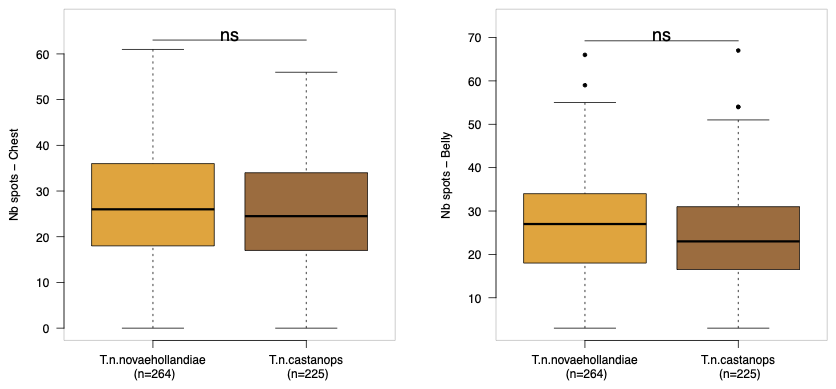

Figure S1 – Boxplots showing the number of eumelanic black spots on the breast (left panel) and belly (right panel) of *Tyto novaehollandiae novaehollandiae* and *T. n. castanops*. Number between brackets report sample sizes. No significant (ns) differences were detected between taxa for either trait.

Welch Two Sample t-test

data: as.numeric(PhenoNova$Wing) by PhenoNova$Subspecies.x

t = -8.6593, df = 431.48, p-value < 2.2e-16

alternative hypothesis: true difference in means between group T.n.novaehollandiae and group T.n.castanops is not equal to 0

95 percent confidence interval:

-19.18125 -12.08460

sample estimates:

mean in group T.n.novaehollandiae	mean in group T.n.castanops

320.4723	336.1053

Table S2 – Results of the *t*-test assessing differences in wing length between *Tyto novaehollandiae novaehollandiae* and *Tyto novaehollandiae castanops*.

Welch Two Sample t-test

data: as.numeric(PhenoNova$Bill) by PhenoNova$Subspecies.x

t = -3.7619, df = 419.98, p-value = 0.0001926

alternative hypothesis: true difference in means between group T.n.novaehollandiae and group T.n.castanops is not equal to 0

95 percent confidence interval:

-1.0983142 -0.3444564

sample estimates:

mean in group T.n.novaehollandiae	mean in group T.n.castanops

23.92790	24.64928

Table S3 – Results of the *t*-test assessing differences in bill length between *Tyto novaehollandiae novaehollandiae* and *Tyto novaehollandiae castanops*.

Welch Two Sample t-test

data: as.numeric(PhenoNova$Claw) by PhenoNova$Subspecies.x

t = -2.7271, df = 32.516, p-value = 0.01022

alternative hypothesis: true difference in means between group T.n.novaehollandiae and group T.n.castanops is not equal to 0

95 percent confidence interval:

-3.5380328 -0.5136036

sample estimates:

mean in group T.n.novaehollandiae	mean in group T.n.castanops

21.83818	23.86400

Table S4 – Results of the *t*-test assessing differences in Claw length between *Tyto novaehollandiae novaehollandiae* and *Tyto novaehollandiae castanops*.

Welch Two Sample t-test

data: as.numeric(PhenoNova$Cpoi) by PhenoNova$Subspecies.x

t = 14.972, df = 330.01, p-value < 2.2e-16

alternative hypothesis: true difference in means between group T.n.novaehollandiae and group T.n.castanops is not equal to 0

95 percent confidence interval:

1.719342 2.239508

sample estimates:

mean in group T.n.novaehollandiae	mean in group T.n.castanops

5.109562	3.130137

Table S5 – Results of the *t*-test assessing differences in breast color between *Tyto novaehollandiae novaehollandiae* and *Tyto novaehollandiae castanops*.

Welch Two Sample t-test

data: as.numeric(PhenoNova$Cven) by PhenoNova$Subspecies.x

t = 13.803, df = 441.77, p-value < 2.2e-16

alternative hypothesis: true difference in means between group T.n.novaehollandiae and group T.n.castanops is not equal to 0

95 percent confidence interval:

1.793351 2.388843

sample estimates:

mean in group T.n.novaehollandiae	mean in group T.n.castanops

5.579681	3.488584

Table S6 – Results of the *t*-test assessing differences in belly color between *Tyto novaehollandiae novaehollandiae* and *Tyto novaehollandiae castanops*.

Welch Two Sample t-test

data: as.numeric(PhenoNova$NbPoi) by PhenoNova$Subspecies.x

t = 0.80501, df = 433.92, p-value = 0.4213

alternative hypothesis: true difference in means between group T.n.novaehollandiae and group T.n.castanops is not equal to 0

95 percent confidence interval:

-1.307025 3.120416

sample estimates:

mean in group T.n.novaehollandiae	mean in group T.n.castanops

26.96610	26.05941

Table S7 – Results of the *t*-test assessing differences in number of spots on the breast between *Tyto novaehollandiae novaehollandiae* and *Tyto novaehollandiae castanops*.

Welch Two Sample t-test

data: as.numeric(PhenoNova$NbVen) by PhenoNova$Subspecies.x

t = 1.8836, df = 459.95, p-value = 0.06024

alternative hypothesis: true difference in means between group T.n.novaehollandiae and group T.n.castanops is not equal to 0

95 percent confidence interval:

-0.08491763 4.01028577

sample estimates:

mean in group T.n.novaehollandiae	mean in group T.n.castanops

26.69880	24.73611

Table S8 – Results of the *t*-test assessing differences in number of spots on the belly between *Tyto novaehollandiae novaehollandiae* and *Tyto novaehollandiae castanops*.

### Ecological differentiation

Table S9 – Records of *Tyto novaehollandiae* from the Global Biodiversity Information Facility (GBIF). GBIF.org. Occurrence Download, accessed 30 October 2025. https://doi.org/10.15468/dl.gwpumg

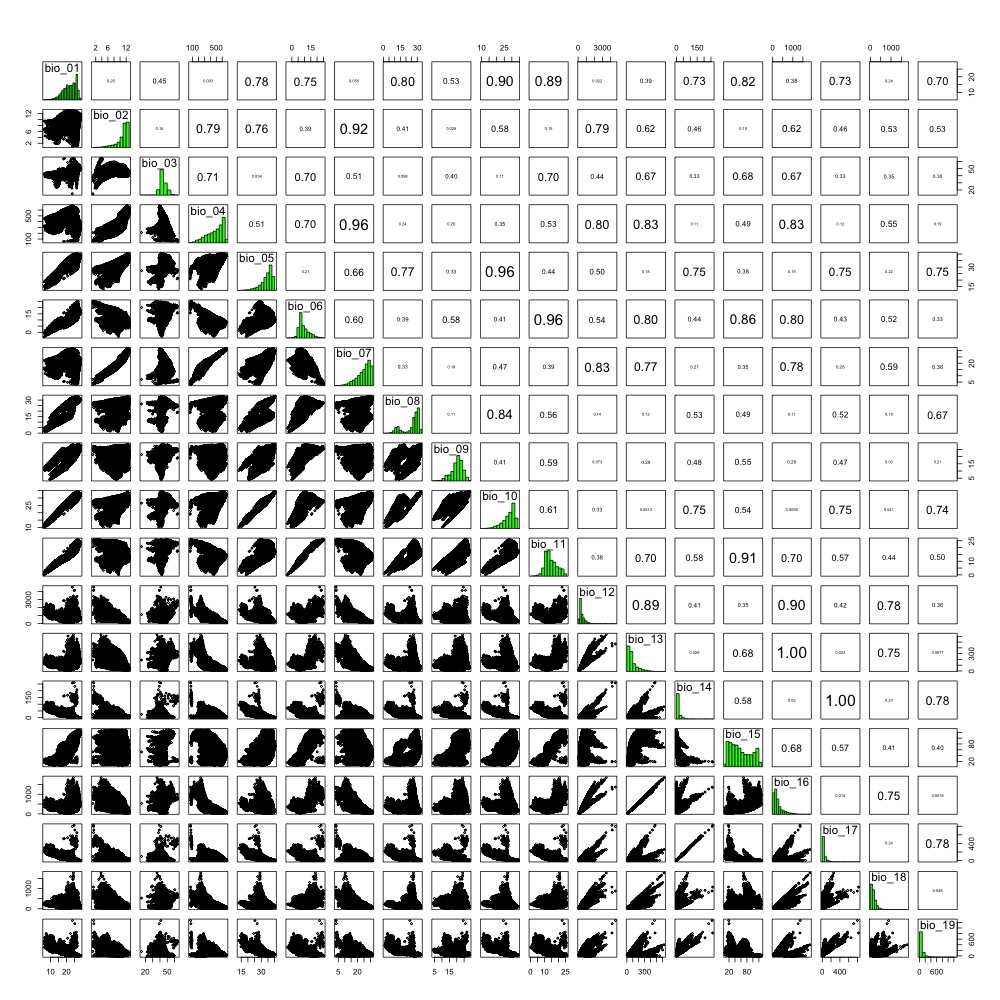

Figure S2 – Correlation between environmental variables across Australia and Tasmania.

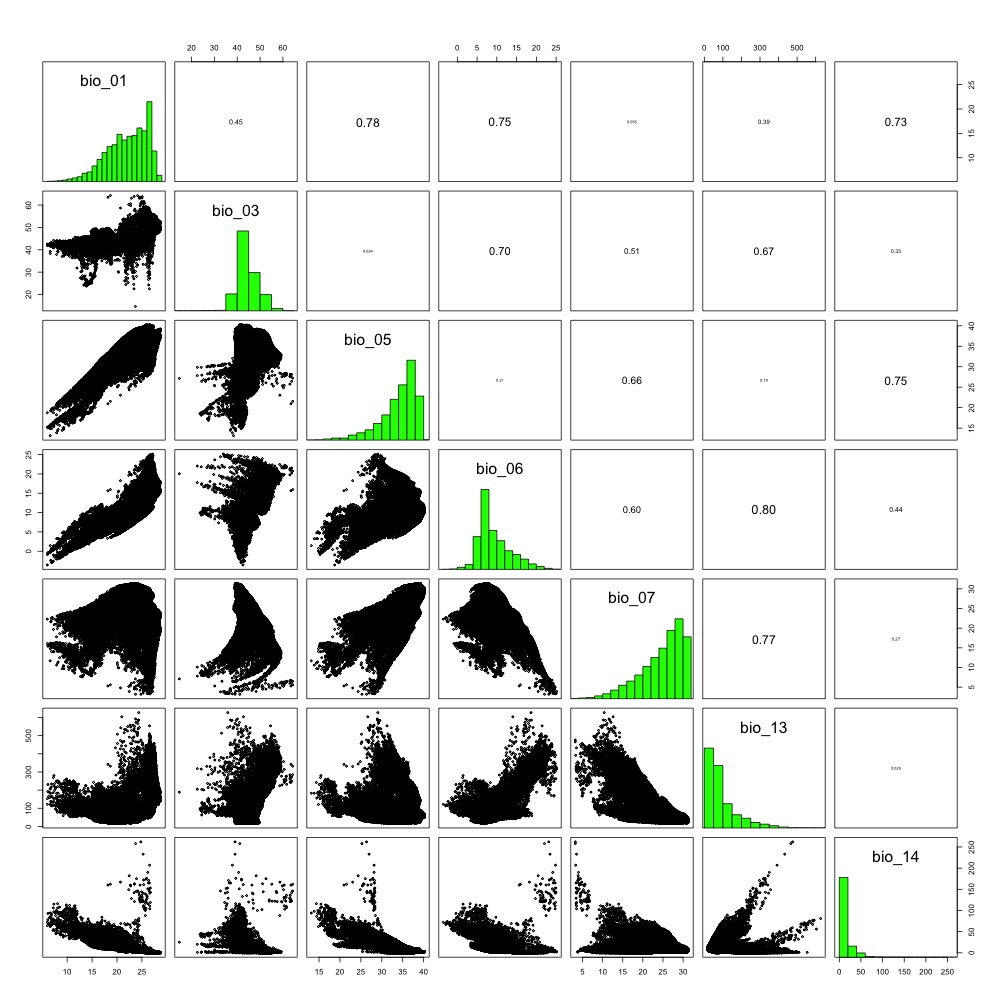

Figure S3 – Correlation between environmental variables across Australia and Tasmania for the set of variables retained for the OMI analysis.

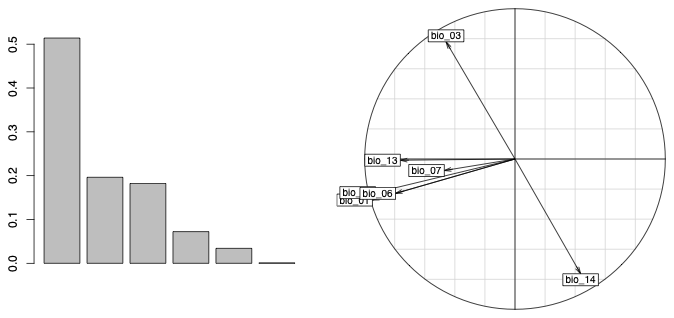

Figure S4 – Principal component analysis (PCA) of climatic variables across the study area. The left panel shows the percentage of total variance explained by each principal component. The right panel displays the loadings of environmental variables on the first two PCA axes (PC1 and PC2), illustrating their relative contribution to the observed climatic variation.

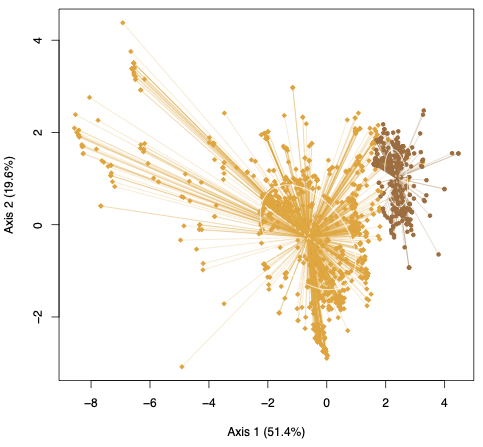

Figure S5 – Distribution of occurrence records in the principal component analysis (PCA) space of climatic variables across the study area. Each point represents an individual occurrence record projected onto the first two PCA axes. Orange diamonds indicate records of *T. n. novaehollandiae* and brown circles indicate records of *T. n. castanops*. Shaded areas were excluded from the analysis.

class: krandtest lightkrandtest

Monte-Carlo tests

Call: as.krandtest(sim = t(sim), obs = obs)

Number of tests: 3

Adjustment method for multiple comparisons: none

Permutation number: 999

Test	Obs Std.Obs	Alter Pvalue

1 environmentAU 0.5235719 767.9663 greater **0.001**

2 environmentTA 7.1683946 866.0488 greater **0.001**

3	OMI.mean 3.8459832 921.2910 greater **0.001**

Table S10 – Results of the Outlying Mean Index (OMI) approach conducted on the occurrences across the study area. Significant results are highlighted in bold.

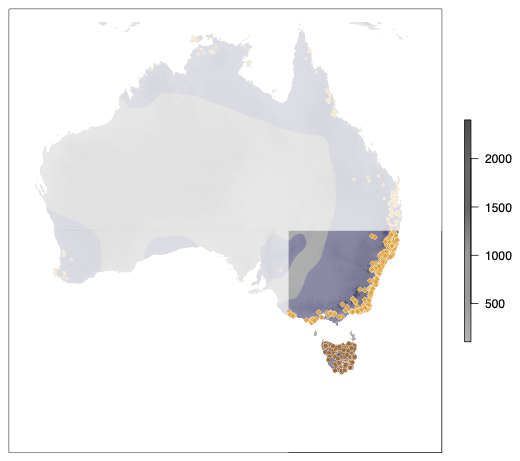

Figure S6 – Restricted study area used for ecological niche analyses. To reduce the influence of the broad climatic heterogeneity encompassed by mainland Australia, analyses were repeated using only occurrence records from southwestern Australia and Tasmania. The dark blue area represents the distribution of *Tyto novaehollandiae*. Orange diamonds indicate records of *T. n. novaehollandiae* and brown circles indicate records of *T. n. castanops*. Shaded areas were excluded from the analysis.

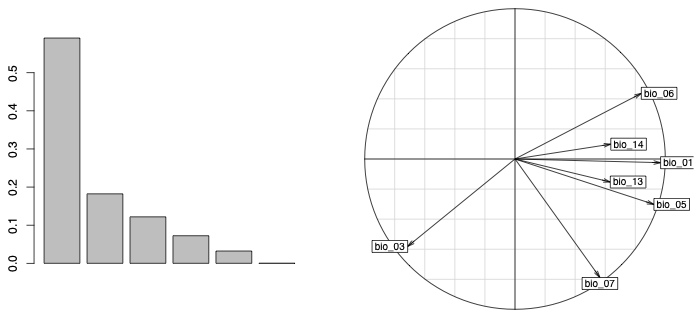

Figure S7 – Principal component analysis (PCA) of climatic variables in the restricted study area. The left panel shows the percentage of total variance explained by each principal component. The right panel displays the loadings of environmental variables on the first two PCA axes (PC1 and PC2), illustrating their relative contribution to the observed climatic variation.

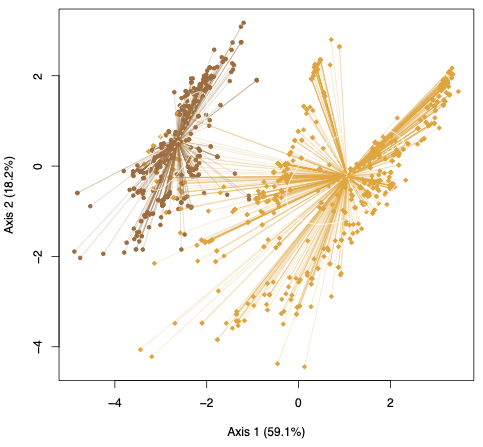

Figure S8 – Distribution of occurrence records in the principal component analysis (PCA) space of climatic variables across the restricted study area. Each point represents an individual occurrence record projected onto the first two PCA axes. Orange diamonds indicate records of *T. n. novaehollandiae* and brown circles indicate records of *T. n. castanops*. Shaded areas were excluded from the analysis.

class: krandtest lightkrandtest

Monte-Carlo tests

Call: as.krandtest(sim = t(sim), obs = obs)

Number of tests: 3

Adjustment method for multiple comparisons: none

Permutation number: 999

Test	Obs	Std.Obs	Alter Pvalue

1 environmentAU 1.167696 858.8700 greater **0.001**

2 environmentTA 7.271916 886.5544 greater **0.001**

3	OMI.mean 4.219806 1012.8784 greater **0.001**

Table S11 – Results of the Outlying Mean Index (OMI) approach conducted on the occurrences across the restricted study area. Significant results are highlighted in bold.

### Genetic differentiation

Table S12 – Description of the museum samples of *T. n. novaehollandiae* and *T. n. castanops* used for the genomic analysis.

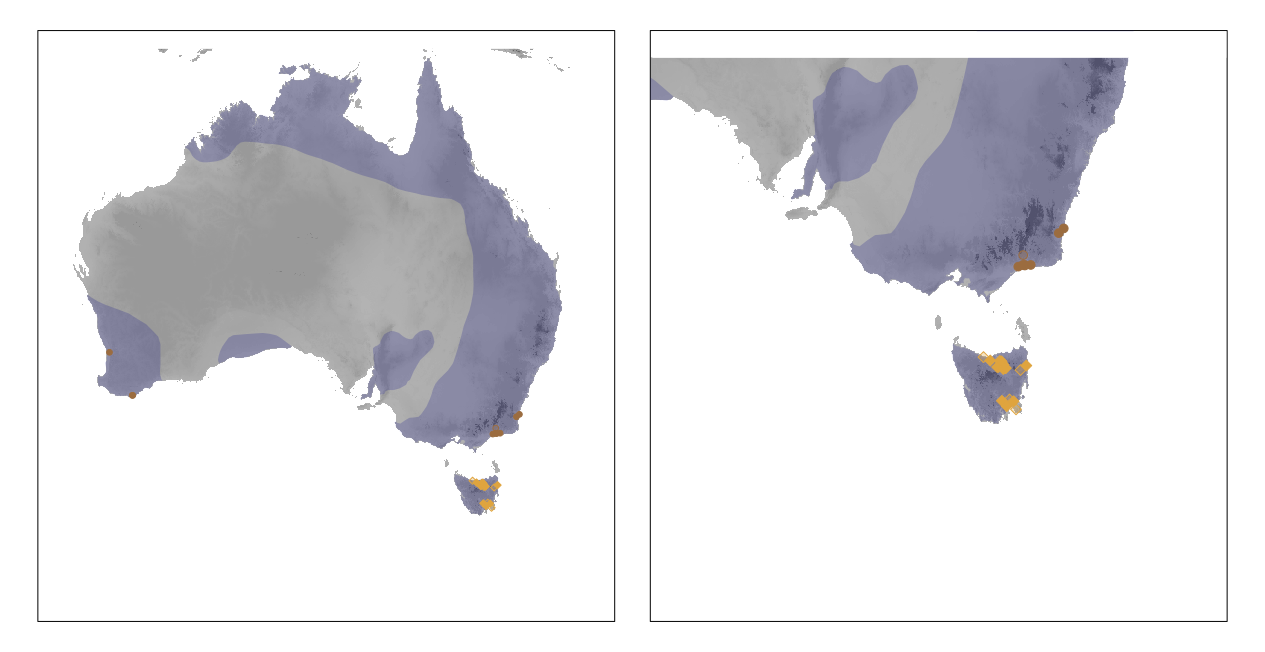

Figure S9 - Geographic distribution of samples sequenced in this study across Australia and Tasmania (left panel) and a zoomed view of southeastern Australia (right panel). The dark blue area represents the distribution of Tyto novaehollandiae. Orange diamonds indicate records of T. n. novaehollandiae, whereas brown circles indicate records of T. n. castanops. Symbols with light-coloured fills represent individuals that were excluded during sample processing and filtering (see the Materials and Methods section for details).

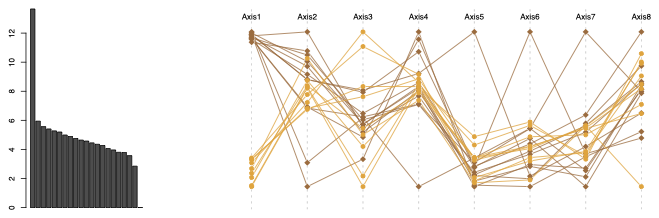

Figure S10 – Principal Component analysis of the genomic data. Screeplot of the 8 fist axes of the PCA with all the individuals (left). Position of the individuals on the 8 first axes of the PCA (right). Orange diamonds indicate records of *T. n. novaehollandiae* and brown circles indicate records of *T. n. castanops*. Shaded areas were excluded from the analysis.

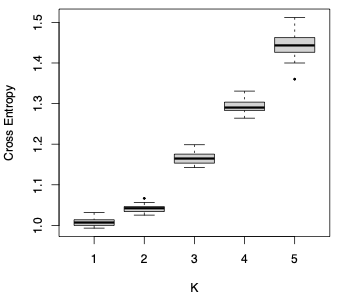

Figure S11 – Values of the cross-entropy criterion for 25 sNMF runs per K. The number of clusters ranged from 1 to 5.

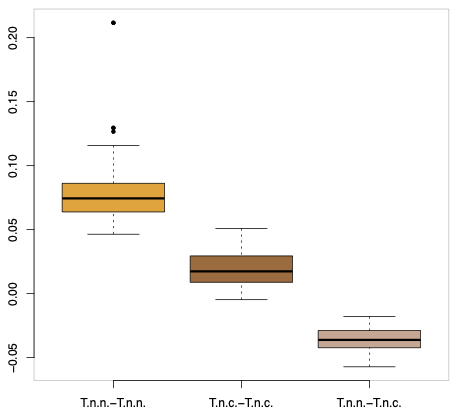

Figure S12 – Boxplot of pairwise kinship coefficients within and between populations. Boxes represent the distribution of kinship coefficients for all pairs of individuals belonging to the same population or to different populations. *T.n.n.* stands for *T. n. novaehollandiae* and *T.n.c.* for *T. n. castanops*.

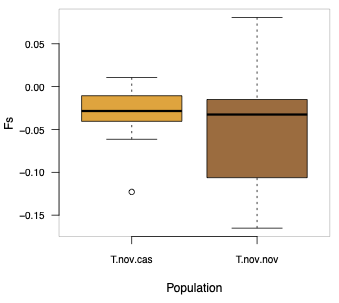

Figure S13 – Boxplot of the inbreeding coefficient estimated using the allele sharing approach (*F*_S_) for each population. Boxes represent the distribution of individual *F*_S_ values within populations, with higher values indicating greater genomic inbreeding.

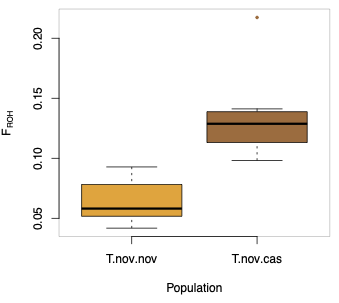

Figure S14 – Boxplot of the inbreeding coefficient estimated from runs of homozygosity (*F*_ROH_) for each population. Boxes represent the distribution of individual *F*_ROH_ values within populations, with higher values indicating greater genomic inbreeding.

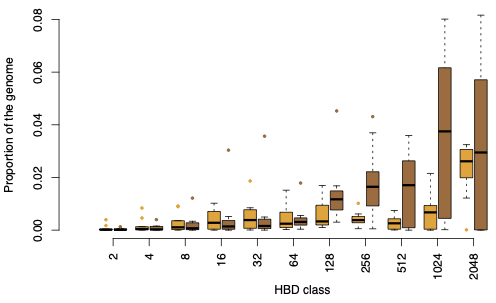

Figure S15 – Boxplot showing the proportion of the genome assigned to different Homozygosity-by-descent (HBD) classes for individuals from *Tyto novaehollandiae novaehollandiae* (orange boxes) and *Tyto novaehollandiae castanops* (brown boxes). HBD classes approximately correspond to different coalescence times, with lower classes representing more recent common ancestry and higher classes representing older ancestry. Boxes represent the distribution of the genomic proportion assigned to each HBD class within each population.

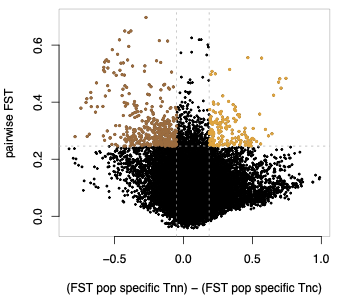

Figure S16 – Distribution of genomic windows according to pairwise *F*_ST_ and the difference in population-specific *F*_ST_ between *Tyto novaehollandiae novaehollandiae* and *T. n. castanops*. The x-axis represents the difference in population-specific *F*_ST_, whereas the y-axis represents pairwise *F*_ST_. Windows showing both elevated pairwise differentiation and an excess of population-specific *F*_ST_ in one of the two populations were classified as candidate differentiated regions and are highlighted.

Table S13 – List of genes overlapping genomic windows identified as significant in *Tyto novaehollandiae novaehollandiae*. Significant windows were defined as exhibiting both elevated pairwise differentiation and an excess of population-specific *F*_ST_ in *T. n. novaehollandiae*.

Table S14 – List of genes overlapping genomic windows identified as significant in *Tyto novaehollandiae novaehollandiae*. Significant windows were defined as exhibiting both elevated pairwise differentiation and an excess of population-specific *F*_ST_ in *T. n. novaehollandiae*.
